# MaizeGDB Phylostrata Tool: Exploring evolutionary origins of maize proteins

**DOI:** 10.64898/2025.12.19.695500

**Authors:** Laura E. Tibbs-Cortes, Oliva C. Haley, John Portwood, Ethalinda Cannon, Margaret Woodhouse, Carson Andorf

## Abstract

**Motivation:** Phylostratigraphic analysis identifies the evolutionary origins and level of conservation of proteins, facilitating research in evolutionary biology and comparative genomics.

**Results:** We developed the MaizeGDB Phylostrata Tool, a custom web application that enables users to explore the evolutionary origins of maize proteins. This tool features interactive visualizations and detailed gene pages incorporating subcellular localization, Gene Ontology (GO) terms, and other resources for homologs. Full-proteome downloads are available for 26 maize inbreds (B73 and the NAM founders). We also provide an updated version of the ‘phylostratr’ R package that makes it more robust to taxonomic updates, as well as example scripts for phylostratigraphic analysis and webtool development for the use of researchers and curators of other species.

**Availability and Implementation:** The MaizeGDB Phylostrata Tool is freely available at phylostrata.maizegdb.org. Scripts used for the analysis and web tool are available at https://github.com/LTibbs/PhylostrataWebtool.

## 1 Introduction

Novel gene evolution is ongoing, giving rise to variation that becomes material for natural or artificial selection. Such genes may originate *de novo* from non-coding sequences, or they may arise from ancestral genes through divergence after duplication, gene fusion or fission, or transposition. Some of these novel genes are quickly lost, while others become founders of new protein families (Domazet-Lošo, Brajković and Tautz 2007; Arendsee, Li and Wurtele 2014; Oss and Carvunis 2019; James *et al*. 2021).

Phylostratigraphy is the process of determining the evolutionary origin of genes and the proteins they encode. This is accomplished by searching for homologous proteins in increasingly broad phylogenetic groups (phylostrata), i.e. subspecies, species, genus, etc. Each protein is assigned to the most basal phylostratum in which a homolog is found, indicating the clade in which the founder gene arose (Domazet-Lošo, Brajković and Tautz 2007; Arendsee *et al*. 2019). For example, a protein may have homologs across cellular organisms (most conserved), it may be species-specific (least conserved), or it may fall somewhere in between (e.g., a protein that arose in grasses).

Phylostratigraphy has many applications in evolutionary biology and comparative genomics. Previous studies have investigated the evolutionary origins of: germ layers and tissue types in *Drosophila melanogaster* (Domazet-Lošo, Brajković and Tautz 2007); regulatory networks in *Populus* xylem (Huang *et al*. 2025); and genes involved in stress response in maize (Hayford *et al*. 2024), *Arabidopsis thaliana* (Mustafin *et al*. 2019), and *Nicotiana attenuata* (Durrant *et al*. 2017). Phylostratigraphy has also been used to identify cases of horizontal gene transfer or gene loss (Arendsee *et al*. 2019), orphan genes (Arendsee, Li and Wurtele 2014), and shifts in protein characteristics over evolutionary time and across taxa (Arendsee, Li and Wurtele 2014; James *et al*. 2021).

The origin of phylostratigraphic analysis coincided with the increasing availability of whole-genome sequencing, enabling the first phylostratigraphic analysis in *Drosophila melanogaster*. This analysis used BlastP to compare protein sequences to all sequences relevant to Drosophila phylogeny in the NCBI non-redundant database, which at the time was only 2.8 million sequences, including 1.4 million sequences from prokaryotes (Domazet-Lošo, Brajković and Tautz 2007).

The concept of building an efficient tree rather than using all available sequences emerged as more genomes were sequenced. In 2020, 150 million non-redundant sequences were available for prokaryotes alone in the NCBI database (Li *et al*. 2021), and comparing proteins to all sequences in NCBI increased computational burden and bias towards highly-sampled strata. In addition, including low-quality proteomes made it difficult to distinguish between a true negative (a given proteome does not include a homolog) and a false negative (a homolog was not found because the proteome is incomplete). The ‘phylostratr’ package, released in 2019, addressed these challenges by building a pruned, diverse clade tree from the UniProt Proteome collection using customizable weights for proteome selection (Arendsee *et al*. 2019).

Here, we provide a custom phylostratigraphic web application available at MaizeGDB with interactive visualization, easy access to resources from homologs in well-studied species as well as protein alignments from these species on MaizeGDB genome browsers, and full-proteome results downloads for the founders of the Maize Nested Association Mapping (NAM) population. We conducted this phylostratigraphic analysis using a version of the ‘phylostratr’ package that we updated to be robust to ongoing revisions in taxonomy. We offer this updated version of ‘phylostratr’, along with sample code for phylostratigraphy and web tool construction, to enable easy phylostratigraphic analysis and user-friendly web tool development in other species.

## 2 Materials and Methods

### 2.1 Updates to phylostratr package

We added two new functions to ‘phylostratr’ (https://github.com/arendsee/phylostratr) to accommodate ongoing taxonomic updates in UniProt, as well as to add more options for customization: ‘generate_prokaryote_tree()’and ‘use_custom_prokaryote_tree()’. The function ‘generate_prokaryote_tree()’automatically accesses the current UniProt taxonomy and uses it to generate a new prokaryote tree based on the ‘diverse_subtree()’ function from the original phylostratr package, rather than using a static tree based on a previous version of the taxonomy. Users can customize these trees by providing the desired number of taxa, taxa weights, and taxa to exclude. We also provide example code for users to download proteome quality information from UniProt (BUSCO and annotation scores) and to incorporate these into weights. Similarly, we worked with phylostratr author Zeb Arendsee (Arendsee *et al*. 2019), to add options to use weights and to exclude taxa to the existing function ‘uniprot_sample_prokaryotes()’. Custom prokaryote trees can then be added to the analysis by using ‘use_custom_prokaryote_tree()’in place of the original ‘use_recommended_prokaryotes()’ function (see vignette at https://github.com/LTibbs/phylostratr/blob/master/vignettes/new_prokaryotes.Rmd).

### 2.2 Phylostratr analysis

We built a custom tree with weights based on BUSCO scores, UniProt annotation scores, and whether organisms were crop and/or model species from the current UniProt taxonomy using the updated ‘phylostratr’ package. This tree included 116 prokaryotes (10 archaea and 106 bacteria) and 57 eukaryotes, with a median BUSCO completeness score of 98%. Forty-eight additional proteomes were added from MaizeGDB, including the maize B73v5 and NAM founders (Yu *et al*. 2008; Hufford *et al*. 2021), Chinese Academy of Agricultural Sciences (CAAS) founders (Wang *et al*. 2023), and other members of the Andropogoneae tribe sequenced in the PanAnd project (Stitzer *et al*. 2025). Including subspecies facilitates distinguishing rapidly evolving genes from *de novo* genes (Arendsee *et al*. 2019). The longest protein isoform was used for each gene model. Blastp from Diamond version 2.1.11 (Buchfink, Reuter and Drost 2021; https://github.com/bbuchfink/diamond) was used to speed up proteome alignments (Arun Seetharam, Purdue University, personal communication). Phylostratr was then run for B73 and for each of the NAM founders.

### 2.3 MaizeGDB Phylostrata Tool

In the MaizeGDB Phylostrata Tool (https://phylostrata.maizegdb.org/) hosted at MaizeGDB (Woodhouse *et al*. 2021a), users can search up to 100 B73v5 (Zm00001eb.1) gene model IDs simultaneously for interactive visualizations of evolutionary conservation as well as detailed gene pages. Additionally, users can download full-proteome results for B73 and the NAM founders from the tool’s Downloads page.

#### 2.3.1 Search page and results

Users begin by entering up to 100 B73v5 gene IDs into the search box (Fig. 1A), one per line. Results are presented as an interactive visualization, showing the level of evolutionary conservation of each protein (Fig. 1B). The 14 phylostrata are arranged from the most conserved on the left to the least conserved on the right, with an example species for each stratum shown at the top. These example species were chosen based on annotation quality (BUSCO completeness and UniProt annotation scores) with a preference for model organisms or crop species. These example species are therefore expected to have more information available for user follow-up.

**Fig. 1:**
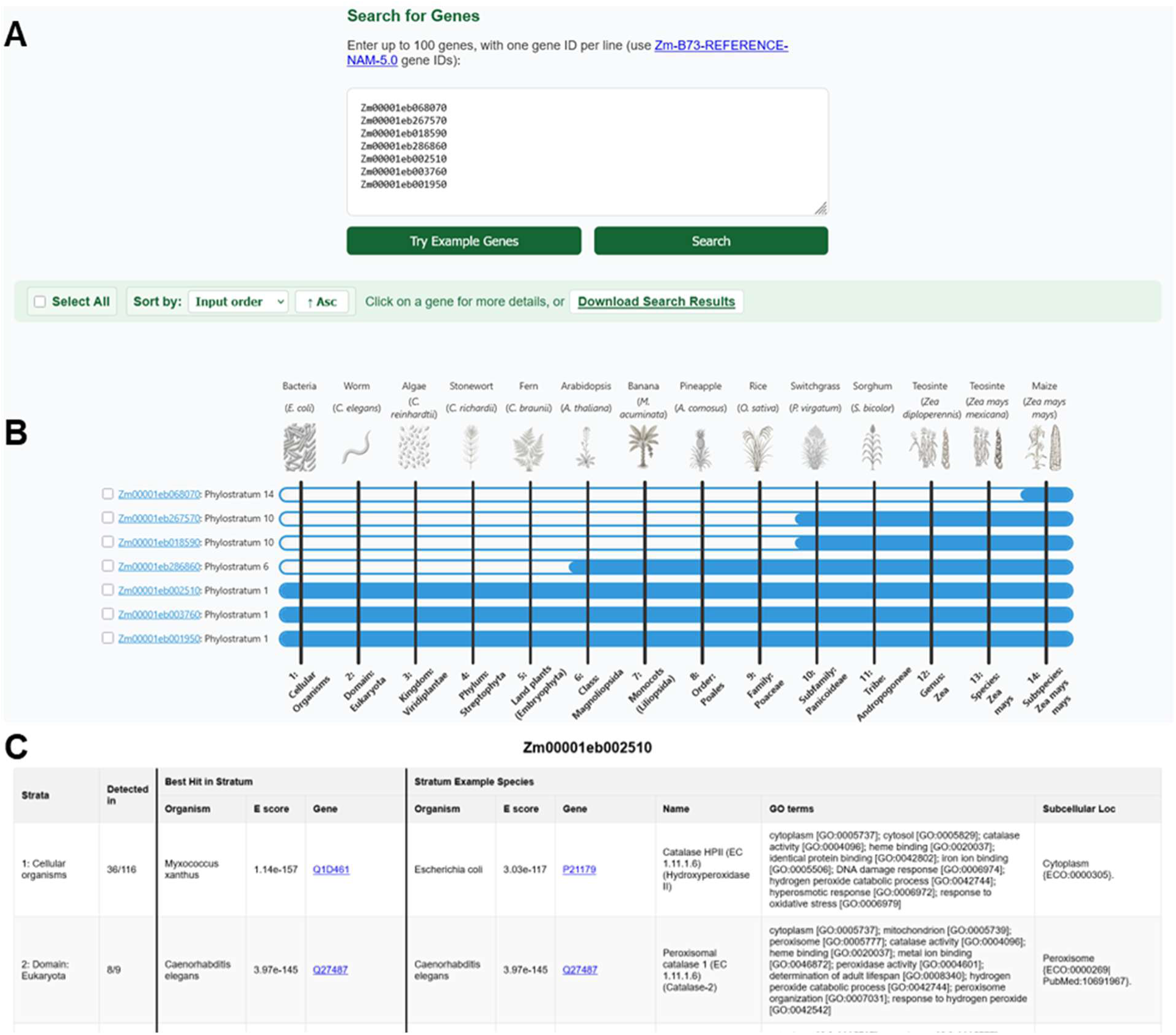
Example use of the MaizeGDB Phylostrata Tool. A) Example genes entered in the search box. B) After clicking “Search”, a result image displays the phylostratum of requested genes in text on the left side of the and visually by the fill of the corresponding blue bar. Results can be selected with checkboxes, sorted, and downloaded. C) Clicking on a gene name brings the user to its Details page, showing details of hits in other species across phylostrata. Here, the top of the Details page for the conserved Zm00001eb002510 (cat2) is shown (see Fig. S6).

The search results display one row for each protein, consisting of a text label including the gene model name and its phylostratum assignment, as well as a blue bar that visually indicates the phylostratum. From right to left, the bar is filled to its phylostratum level for a given protein; a more filled bar indicates a more conserved protein. Users can sort this visualization in ascending or descending order by input, phylostratum, or alphabetically, enabling easy visual comparison of proteins. Checkboxes enable users to select genes of interest. Finally, the customized image can be downloaded as a PNG.

#### 2.3.2 Details pages

Each gene model name in the search results includes a hyperlink (Fig. 1B) to a Details page for its encoded protein (Fig. 1C). Each row on the Details page represents one phylostratum, displaying organism name, E-value, and gene ID for the best hit overall (the highest-scoring homolog found in any species in the current phylostratum) and the best hit in the example species (the highest-scoring homolog found in the example species of the current phylostratum). Both the best hits and example hits are hyperlinked to associated UniProt or MaizeGDB entries. Example species are the same as those used in the results visualization.

Gene name, Gene Ontology (GO) terms, and subcellular localization are also shown for the best hit in the example species. These data were obtained from UniProt when available, or predicted using EnTAP v1.4.0 (Hart *et al*. 2020) and DeepLoc 2.1 (Nielsen 2025). For EnTAP, the ‘uniprot_sprot’ and ‘refseq_plant’ DIAMOND databases were used (https://treegenesdb.org/FTP/EnTAP/latest/databases/, version 2025-01-11). Insect, fungi, and bacterial contaminants were removed (‘contam=insecta,fungi,bacterià). In DeepLoc, the fast model was used for proteins shorter than 1022 residues (‘-m Fast’) and the accurate model for longer proteins (‘-m Accuratè). The Details page also shows the proportion of species in each phylostratum in which homologs of the query protein were found. Users can download all or selected Details pages from search results as html or txt files.

#### 2.3.3 Full-proteome downloads

From the Downloads page, users can download full proteome results for B73 and the maize NAM founders: phylostrata_results files, which contain only the gene ID and corresponding phylostratum; and phylostrata_details files, which contain all information from the Details pages.

#### 2.3.4 Phylostrata results incorporated in MaizeGDB gene pages

Single-gene results images were generated for the longest isoform of each B73v5 gene model and added to the associated gene model pages at MaizeGDB. Links to the relevant Details page and to the About page of the Phylostrata Tool are provided (Fig. S1).

### 2.4 Generalized phylostrata webtool

We provide generalized code to create versions of this web tool for other species. The GitHub repository (https://github.com/LTibbs/PhylostrataWebtool) contains scripts to process ‘phylostratr’ output and UniProt downloads into JSON files required for the web tool visualization, as well as code to build the web tool itself. Sections that require customization for different species are marked with “NOTE” in the code.

### 2.5 Additional analyses of phylostrata results

Miniprot v0.18-r281 (Li 2023) with the filtering option ‘--outs 0.9’ was used to align the proteomes of 11 example species (those without genomes at MaizeGDB) to the maize NAM and PanAnd genomes. The GFF files from these alignments were added as tracks on the respective MaizeGDB browsers, enabling users to visualize alignments among homologs (Fig. S2).

RNA-seq data from 241 B73 tissues across 28 projects were downloaded from MaizeGDB’s qTeller (Woodhouse *et al*. 2021b). A list of differentially expressed genes (DEGs) in B73 under abiotic, biotic, or both stress types was obtained from (Hayford *et al*. 2024). GO term enrichment was calculated using ‘enricher()’ from the R package ‘clusterProfiler’ with default FDR correction for multiple testing (Wu *et al*. 2021).

## 3 Results

### 3.1 Usage example of MaizeGDB Phylostrata Tool

Here, we provide an example use of the MaizeGDB Phylostrata Tool with seven maize proteins of varying levels of conservation (Fig. 1).

The least conserved of these example genes is Zm00001eb068070, with significant hits only in *Zea mays mays* (Phylostratum 14, Fig. 1B). Clicking on the gene name takes the user to the associated Details page (Fig. 1C, Fig. S3), which displays GO terms and subcellular localization for homologs in example species. Because Zm00001eb068070 is a maize-specific gene, details for this gene are available only from B73, not other species. This Details page shows that Zm00001eb068070 is associated with the GO terms nucleic acid binding [GO:0003676] and zinc ion binding [GO:0008270], and its predicted subcellular localization is in the cytoplasm or nucleus, providing clues to the potential functions of this species-specific gene (Fig. S3).

Zm00001eb267570 (zein protein, 15kDa15, zp15) and Zm00001eb018590 are both conserved at the subfamily level (Phylostratum 10, Fig. 1B). For zp15, the Details page shows similar GO terms (nutrient reservoir activity) and subcellular localization (extracellular signal peptide, soluble) across example species, indicating likely conservation of function across these species (Fig. S4). Alternatively, in cases such as Zm00001eb018590 where less information is available in maize, the additional information available for other species in the Details page (Fig. S5) can help the user to further investigate this protein’s function. Clicking on hits from NAM or PanAnd genomes on the Details page directs to their genome browsers at MaizeGDB, with miniprot alignments of proteins from the example species displayed for visualization (Fig. S2).

The model organism *Arabidopsis thaliana* represents the class Magnoliopsida (Phylostratum 6) (Fig. 1B). Therefore, users can take advantage of the wealth of information available for Arabidopsis to infer the function of target genes conserved at Phylostratum 6 or earlier. The hyperlinks on the Details page provide easy access to homolog UniProt pages and the information they contain, including expression, interaction, structure, publications and more. For example, at the time of writing, 15 publications are indexed in UniProt for the Arabidopsis homolog of Zm00001eb286860 (tangled1, tan1).

The example genes Zm00001eb002510 (catalase2, cat2), Zm00001eb003760 (lethal leaf spot1, lls1), and Zm00001eb001950 (PBA1 homolog1, pba1) are conserved across cellular organisms (Phylostratum 1). According to UniProt annotations displayed in the Details page, cat2 is a conserved catalase found in the peroxisome across phylostrata (Fig. S6). However, even broad conservation does not guarantee identical function, and further investigation is always needed to confirm function across species. For example, lls1 is an oxygenase found throughout cellular organisms, but has the GO term defense response to bacterium [GO:0042742] beginning in Phylostratum 6 (class: Magnoliopsida) (Fig. S7). This suggests a potential new function of this old gene; in maize, this gene influences resistance to Curvalaria leaf spot (Li *et al*. 2022).

Finally, the Details page for pba1 (Fig. S8) shows that it is assigned to Phylostratum 1 because it is present in the cyanobacterium *Lusitaniella coriacea*, and is found throughout Viridiplantae, but it is absent in all of the non-photosynthetic eukaryotes (Phylostratum 2). For pba1 and its homologs, GO terms include chloroplast thylakoid membrane [GO:0009535] and protein names include cyanobacterial aminoacyl-tRNA synthetase (CAAD) domain-containing protein. Therefore, this “skipped” stratum for pba1 provides a likely example of horizontal gene transfer, reflecting the evolutionary origin of the chloroplast as a cyanobacterium (Martin and Kowallik 1999; Stadnichuk and Kusnetsov 2021).

### 3.2 Phylostratigraphic analysis of diverse maize proteomes

#### 3.2.1 Proteome-wide phylostrata distribution in B73 and the NAM founders

The oldest, most conserved proteins tended to be more common than the youngest, least conserved proteins on a proteome-wide scale (Fig. 2A), with Phylostratum 1 (cellular organisms) accounting for 24.5% of each proteome on average, followed by Phylostratum 2 (domain Eukaryota) at 18.4%. Fewer proteins were assigned as teosinte-specific (0.90% assigned to Phylostratum 13 and 0.95% to Phylostratum 12) than maize-specific (4.4% assigned to Phylostratum 14). This may be due to the larger number of domesticated maize genomes used in this analysis (26 *Zea mays mays*) compared to teosinte (eight total teosinte, including five *Zea mays* and three other *Zea* samples), allowing for better identification of maize-specific genes.

**Fig. 2:**
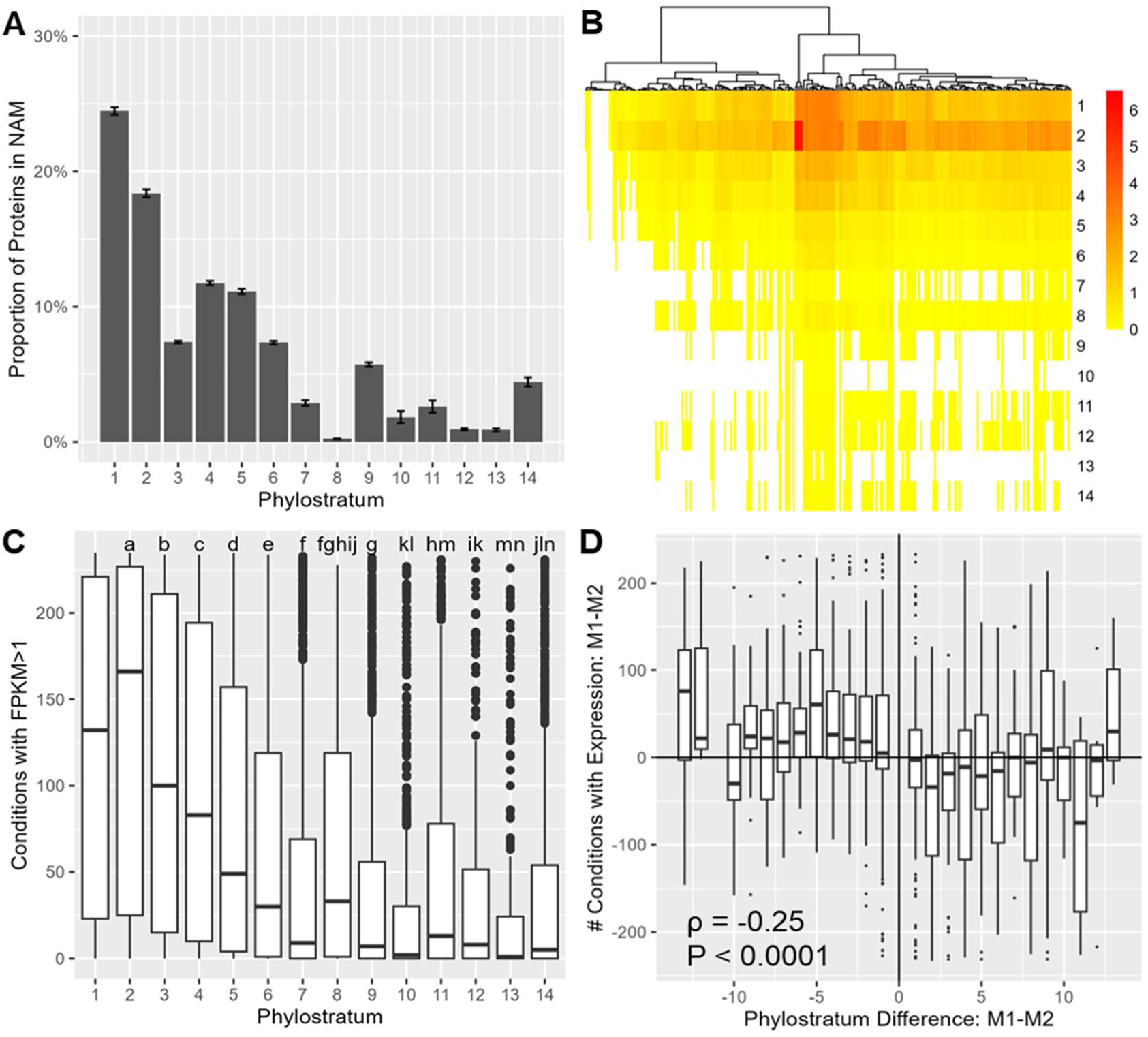
Phylostrata patterns. A) Mean proportion of proteins assigned to each phylostratum in B73 and the other NAM founders; error bars show standard deviation. B) Median expression (log_2_ of FPKM) by phylostratum in 241 B73 tissues. Each column shows one tissue and each row one phylostratum (see Fig. S14). C) Expression breadth across phylostrata. Shared letters indicate phylostrata that are not significantly different using a pairwise Wilcoxon test with FDR correction for multiple testing. D) Expression breadth in homeologs assigned to different subgenomes (see Fig. S12). The left of this panel (M1-M2<0) shows cases where the M1 homeolog is more conserved and the right (M1-M2>0) shows cases where the M2 homeolog is more conserved.

Analyses of the NAM founder accessions have demonstrated that tropical NAM cultivars tend to show greater resistance to certain pathogens than temperate cultivars (Poland *et al*. 2011; Singh *et al*. 2023); previous work also suggested that genes associated with response to biotic stresses tend to be evolutionarily younger than genes involved in response to abiotic stresses (Hayford *et al*. 2024). Therefore, we asked whether tropical NAM lines would show a higher proportion of younger genes than temperate lines, with the expectation that they might be involved in pathogen resistance. However, we found that the proportion of proteins assigned to each phylostratum did not differ significantly across genotype groups (temperate, tropical, mixed, popcorn, and sweetcorn) in the NAM founders (Fig. S9).

#### 3.2.2 Gene length and expression by phylostratum

When analyzing trends in gene model size, we found that more conserved proteins tended to be longer than newer proteins (ρ = -0.43, P < 0.0001) and have more exons (ρ = -0.47, P < 0.0001) (Fig. S10). In B73, the median protein conserved across cellular organisms had 422 residues and six exons, while a median species-specific protein was only 134 residues long with one exon.

Across 241 RNA-seq data sets within B73 spanning a broad range of tissues and conditions (Woodhouse *et al*. 2021b) (https://qteller.maizegdb.org/), expression differed significantly by phylostratum, with higher expression in more conserved phylostrata (*ρ* = -0.76, P < 0.0001) (Fig. 2B, Smedianexpr). The expression of less conserved genes was also more variable as measured by the coefficient of variation (*ρ* = 0.39, P < 0.0001).

Breadth of expression, measured by the number of conditions in the qTeller database with FPKM>1, also varied across phylostrata, with more conserved genes expressed in more conditions (*ρ* = -0.363, P < 0.0001). For example, genes in Phylostratum 1 (most conserved) were expressed in a median of 134 of 241 conditions, while those in Phylostratum 14 (least conserved) were expressed in a median of 7 of 241 conditions (Fig. 2C). Overall, 97.6% of genes were expressed with FPKM>1 in at least one condition, including 93.4% of maize-specific genes (Phylostratum 14).

Finally, we examined published DEGs under various stress conditions (abiotic, biotic, or both) (Hayford et al. 2024). We found that phylostratum differed significantly by stress type (P < 0.0001), with DEGs under abiotic stress tending to be less conserved (mean Phylostratum 3.58) than those under biotic stress (3.00), and DEGs in both conditions in between (3.24) (Fig. S11). This was apparently in contrast to the original publication of these DEGs, which reported that DEGs associated with biotic stress tended to be younger than those associated with abiotic stress (Hayford et al. 2024). However, we found that this difference was due to the previous study reporting phylostrata only for approximately 10% of DEGs, focusing on the subset of DEGs assigned to high-ranked clusters by MCODE (Hayford et al. 2024). When all DEGs were included, their results also showed that DEGs under biotic stress tended to be most conserved, followed by DEGs under both conditions, and finally those under abiotic stress, agreeing with our observed pattern (Rita Hayford, Delaware State University, personal communication). This suggests that, on a genome-wide scale in maize, genes responding to biotic stress tend to have a more ancient evolutionary origin than those responding to abiotic stress.

#### 3.2.3 Phylostrata of B73 subgenomes

Similar relationships of phylostratum with protein length as well as with expression level and breadth were found when comparing homeologs across the two maize subgenomes. The maize ancestor underwent a whole-genome duplication event 10-15 MYA (Stitzer *et al*. 2025), followed by higher rates of gene deletion and loss of function within Subgenome 2 (M2) than Subgenome 1 (M1) (Schnable, Springer and Freeling 2011; Hufford *et al*. 2021; Stitzer *et al*. 2025).

We found that the majority of homeologs were assigned to the same phylostratum (80%, 4216 of 5258 pairs) or within one phylostratum of each other (87%, 4590 pairs), but where there were differences in phylostratum assignments, the homeolog from M1 tended to be assigned as more conserved. M1 also tended to have the longer homeologous protein by mean 7.4 residues and 0.07 exons, as well as to be more highly expressed by 1.0 FPKM and expressed (at FPKM>1) in 8.0 more conditions. However, among the 9.5% of protein pairs where M2 was assigned as more conserved, these patterns were reversed: the protein from M2 tended to be longer (by mean 73.5 residues and 0.95 exons), and to be more highly and widely expressed (by mean 0.5 FPKM and 21.5 tissues), indicating that phylostrata differences between subgenomes largely correspond to differences in length and expression. Among homeologs assigned to different phylostrata, differences in exon number were most correlated with differences in phylostratum (ρ = -0.30, P < 0.0001) followed by those in the number of conditions with expression (ρ = -0.25, P < 0.0001), median expression (ρ = -0.24, P < 0.0001), and protein length (ρ = -0.22, P < 0.0001) (Fig. 2D, Fig. S12).

#### 3.2.4 Subcellular localization and GO terms by phylostratum

Phylostrata also varied significantly by subcellular localization (Kruskal-Wallis rank sum test, P < 0.0001). Proteins located in the chloroplast and plastid were most conserved, while extracellular proteins were the least conserved (Fig. S13). Notably, the majority of proteins localized in the chloroplast were common to cellular organisms (59%) or to Viridiplantae (14%), while very few were common to eukaryotes (3%), reflecting the evolutionary history of the chloroplast as cyanobacteria (Martin and Kowallik 1999; Stadnichuk and Kusnetsov 2021) (Fig. S13).

When analyzing GO term enrichment in each phylostratum (Table S1), we found that 813 GO terms were significantly enriched (FDR-adjusted P < 0.05) in the most conserved phylostratum, with the most significant being ATP hydroloysis activity [GO:0016887], chloroplast stroma [GO:0009570], and peptidyl-tyrosine modification [GO:0018212], aligning with the deep evolutionary origins of these functions. In the younger phylostrata, there were no significantly enriched GO terms in Phylostrata 14 and 12, but two terms (mitochondrial DNA-directed RNA polymerase complex [GO:0034245] and DNA-directed 5’-3’ RNA polymerase activity [GO:0003899]) were enriched in Phylostratum 13 and one term (transposition [GO:0032196]) was enriched in Phylostratum 11.

## 4. Discussion

Several of the phylostratigraphic patterns we identified in maize have been previously reported in other plant species. Our finding that more conserved proteins tend to be more common than younger proteins is consistent with previous work in *Arabidopsis thaliana* (Arendsee, Li and Wurtele 2014; Arendsee *et al*. 2019; Mustafin *et al*. 2019) and *Populus* species (Huang *et al*. 2025), as well as with expectations based on gene origination rates (Ding, Zhou and Wang 2012; Oss and Carvunis 2019) and the evolutionary time represented by each stratum (Domazet-Lošo, Brajković and Tautz 2007). Similarly, increased protein length and/or exon number with protein age has previously been reported in *Arabidopsis thaliana* (Guo 2013; Arensee, Li and Wurtele 2014) and *Populus* species (Huang *et al*. 2025), and higher expression in more conserved phylostrata has been reported in *Arabidopsis thaliana* (Guo 2013). These patterns also align with the general expectation that *de novo* genes encode smaller proteins and have lower expression (Ding, Zhou and Wang 2012; Oss and Carvunis 2019). The consistency of our results in maize with previous phylostratigraphic analyses in two other plant orders suggests that these are likely general trends, while providing support for the efficacy of our updated ‘phylostratr’ package.

Maize and other members of Andropogoneae have a significant history of polyploidy, with at least a third of speciations in this family associated with duplication events (Estep *et al*. 2014; Stitzer *et al*. 2025). In *Zea*, the paleotetraploid ancestor was reduced to a diploid via chromosome reduction, gene loss, and transposon amplification, but more than 5,000 homeologous pairs remain in the genome (Stitzer *et al*. 2025). Of these pairs, we found that a majority (80%) were assigned to the same phylostratum, reflecting the common evolutionary origin of homeologs. However, among pairs with members assigned to different phylostrata, we found that M1 was typically assigned to the older phylostratum (Fig. 2D, Fig. S12). This likely reflects the biased fractionation process in maize, in which genes from M1 were disproportionately likely to be retained. This leaves the M2 copy under reduced selective constraint, facilitating decay, loss, or neofunctionalization to become a novel gene. In fact, we found that members of M2 tend to display characteristics of decayed or novel genes, including reduced expression and smaller protein size (Oss and Carvunis 2019), when compared to their M1 homeologs. However, in cases where the M2 copy was assigned to the older phylostratum, these patterns were reversed, with M2 showing higher expression than M1. This corresponds with previous findings that the more expressed homeolog is more likely to be retained (Schnable, Springer and Freeling 2011), and indicates that, in such cases, the M1 homeolog is likely under reduced selection constraint rather than the M2 copy, resulting in exceptions to the general biased fractionation in maize. These findings provide insight into differential fractionation and the fate of genes after genome-duplication events.

Another notable result was the observed enrichment of GO terms for DNA-directed RNA polymerase and transposition among proteins assigned to Phylostratum 13 (species *Zea mays*) and 11 (tribe Andropogoneae), respectively. This enrichment aligns with the large contribution of transposable elements (TEs) to maize (>85% of base pairs (Stitzer *et al*. 2021)) and Andropogoneae (mean 66% across 8 species (Ramachandran *et al*. 2020)) compared to other plants (mean 47.6% across 67 species (Pedro *et al*. 2021)). Individual TEs have impacted important traits in maize, famously including the insertion of a TE in the regulatory region of the domestication gene *teosinte branched1* (*tb1*) causing increased expression and therefore apical dominance in maize compared to teosinte (Studer *et al*. 2011). TEs in maize can also facilitate gene amplification or be co-opted as gene promoters, shaping gene regulation more broadly (Bubb *et al*. 2025). Enrichment of these GO terms in Andropogoneae and *Zea mays* likely reflects the unusually large impact of TEs in these taxa.

## 5. Conclusion

Through phylostratigraphic analysis of diverse maize accessions, we found that evolutionarily ancient proteins were more common than those with recent origins; however, maize-specific proteins still make up a mean of 4.4% of proteins across the NAM founders (Fig. 2, Fig. S9). We also found that more conserved proteins tended to be longer and to be expressed at higher levels across more conditions, both at a genome-wide scale and when comparing between sub-genomes, providing insight into gene fates after genome duplication. Subcellular localizations and GO terms varied significantly with phylostratum, and genes responding to biotic stresses tended to be more conserved than those responding to abiotic stresses.

By providing our results in a user-friendly web tool that incorporates additional details about homologs, we enable maize genetics researchers to easily investigate the evolutionary origin and conservation of genes of interest; compare and contrast GO terms and subcellular localization of homologs across species; and access protein alignments, publications, and other resources from well-studied species. Our available downloads also facilitate investigation of proteome-wide patterns in B73 and the NAM founders. In addition, our updated tree-building functions make ‘phylostratr’ analyses robust to taxonomic updates. Finally, we have provided scripts to build a customizable web tool for any species, assisting curators and database managers in display and dissemination of phylostratigraphic results. Together, these resources will assist researchers in comparative genomics and gene function prediction across species.

## Supporting information

TableS1_TibbsCortesetal

## Acknowledgements

Funding and Acknowledgments

This research was supported by the U.S. Department of Agriculture, Agricultural Research Service, Project Number 5030-21000-072-00-D through the Corn Insects and Crop Genetics Research Unit in Ames, Iowa. LTC was supported by a postdoctoral fellowship administered by the Oak Ridge Institute for Science and Education (ORISE) through an interagency agreement between the U.S. Department of Energy (DOE) and the U.S. Department of Agriculture (USDA). ORISE is managed by ORAU under DOE contract number DE-SC0014664. This research used resources provided by the SCINet project and/or the AI Center of Excellence of the USDA Agricultural Research Service, ARS project numbers 0201-88888-003-000D and 0201-88888-002-000D.All opinions expressed in this paper are the authors’ and do not necessarily reflect the policies and views of USDA, DOE, or ORAU/ORISE. USDA is an equal opportunity provider and employer.

## Data Availability and Code

All data were obtained from publicly available sources (https://maizegdb.org/, https://www.uniprot.org/). The updated phylostratr package is available at https://github.com/LTibbs/phylostratr (forked from https://github.com/arendsee/phylostratr). Scripts used for the phylostratigraphic analysis and web tool reported in this paper, as well as generalized code for analysing and visualizing other species, can be found at https://github.com/LTibbs/PhylostrataWebtool. Code for subcellular localization and GO term annotation is available at https://github.com/LTibbs/SCINet_tutorials.

## Use of LLM

We used ChatGPT with manual intervention and inspection to generate images of example species for Phylostrata 1 – 11 for the web tool results visualization.

## CRediT Taxonomy

LETC: Conceptualization, Data curation, Formal analysis, Investigation, Methodology, Software, Visualization, Writing -original draft, Writing -review & editing; OCH: Writing -review & editing; JP: Software, Writing -review & editing; EC: Writing -review & editing; MW: Conceptualization, Funding acquisition, Resources, Supervision, Visualization, Writing -review & editing; CA: Conceptualization, Funding acquisition, Resources, Supervision, Visualization, Writing -review & editing

---

**Fig. S1:**
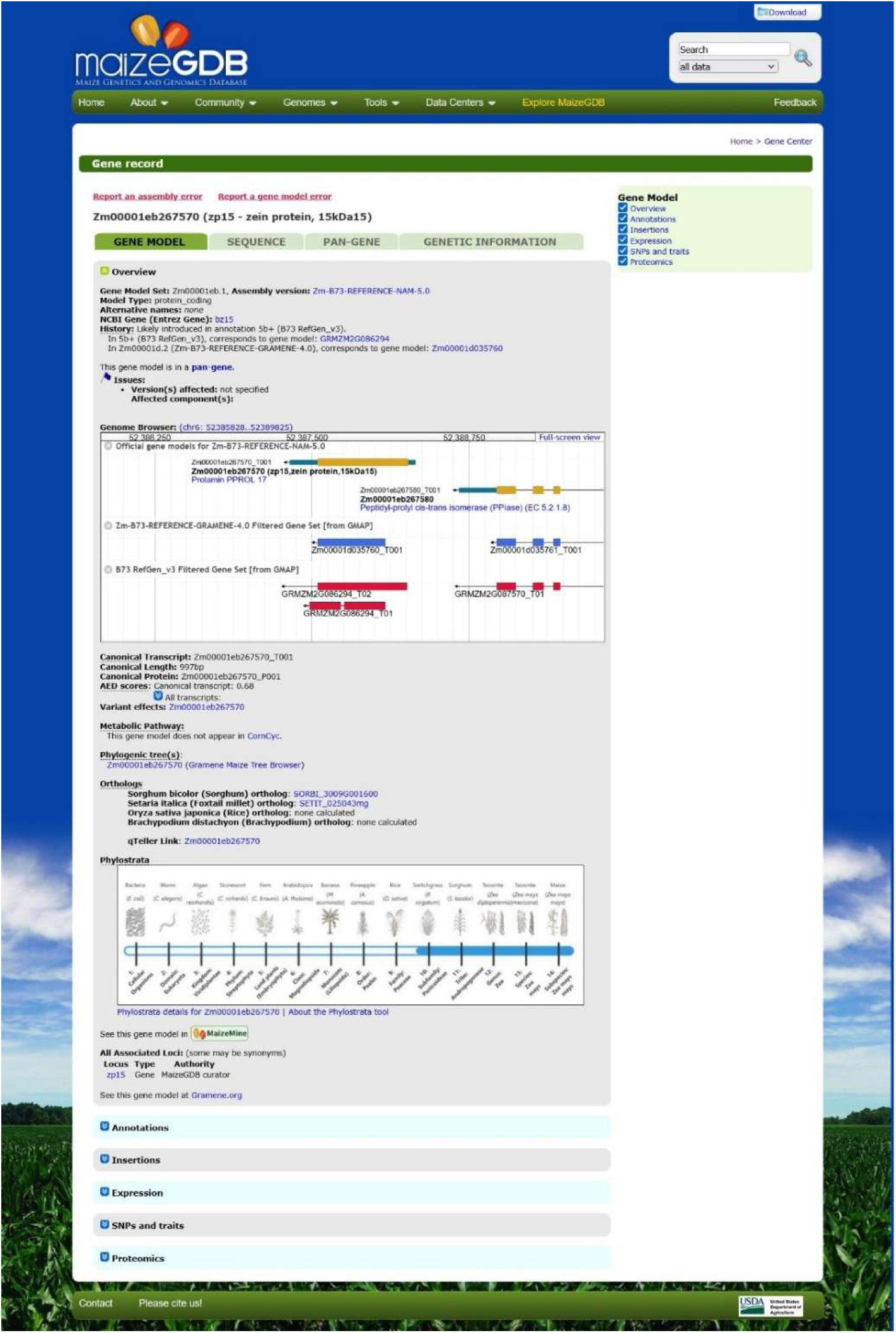
Example gene page showing single-gene phylostrata results image near the bottom of the Gene Model Overview, along with links to the associated Details page and the About page of the Phylostrata Tool.

**Fig. S2:**
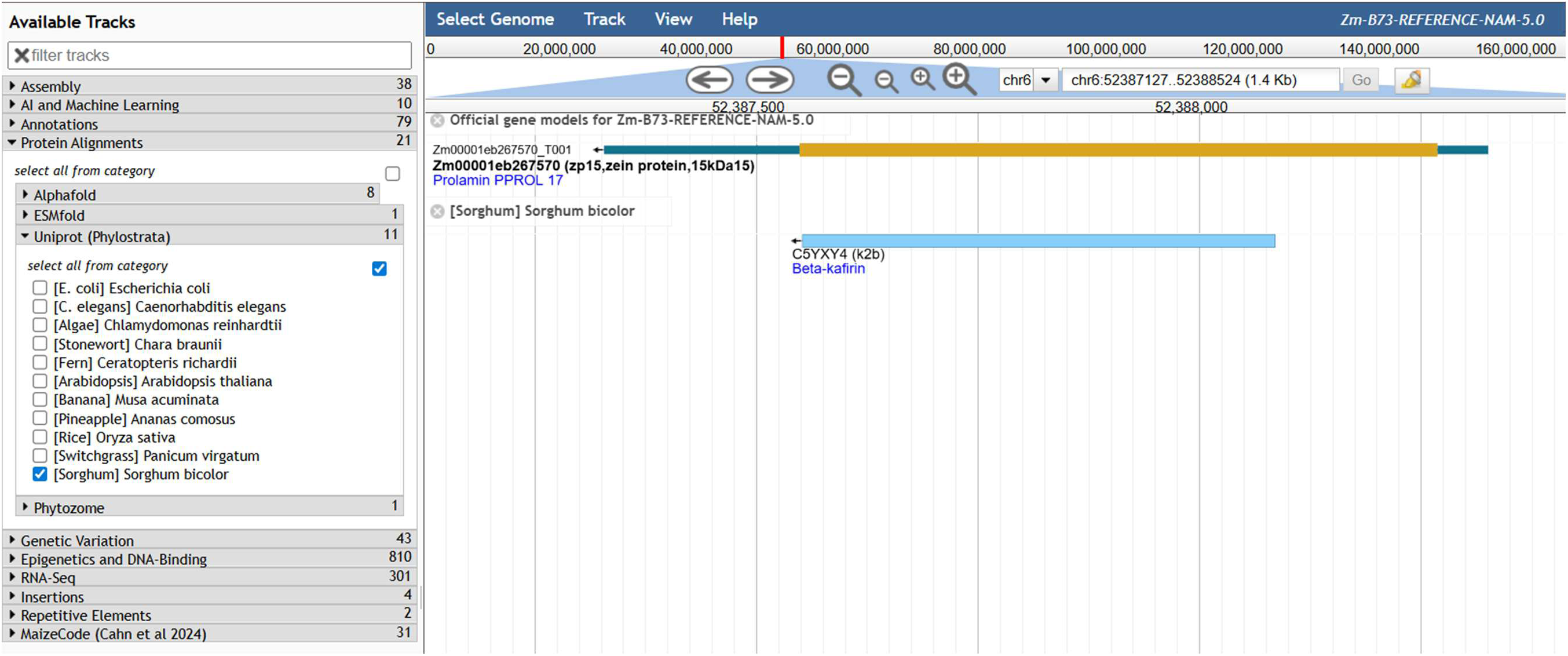
Clicking on a maize or teosinte gene name in a Details page brings the user to the associated browser and shows the miniprot alignment for example species. Here, the alignment of Zm00001eb267570 with sorghum is shown in the B73 genome browser. The alignment tracks can be accessed directly in the browser by clicking the “Uniprot (Phylostrata)” checkboxes under the “Protein Alignments” category.

**Fig. S3:**
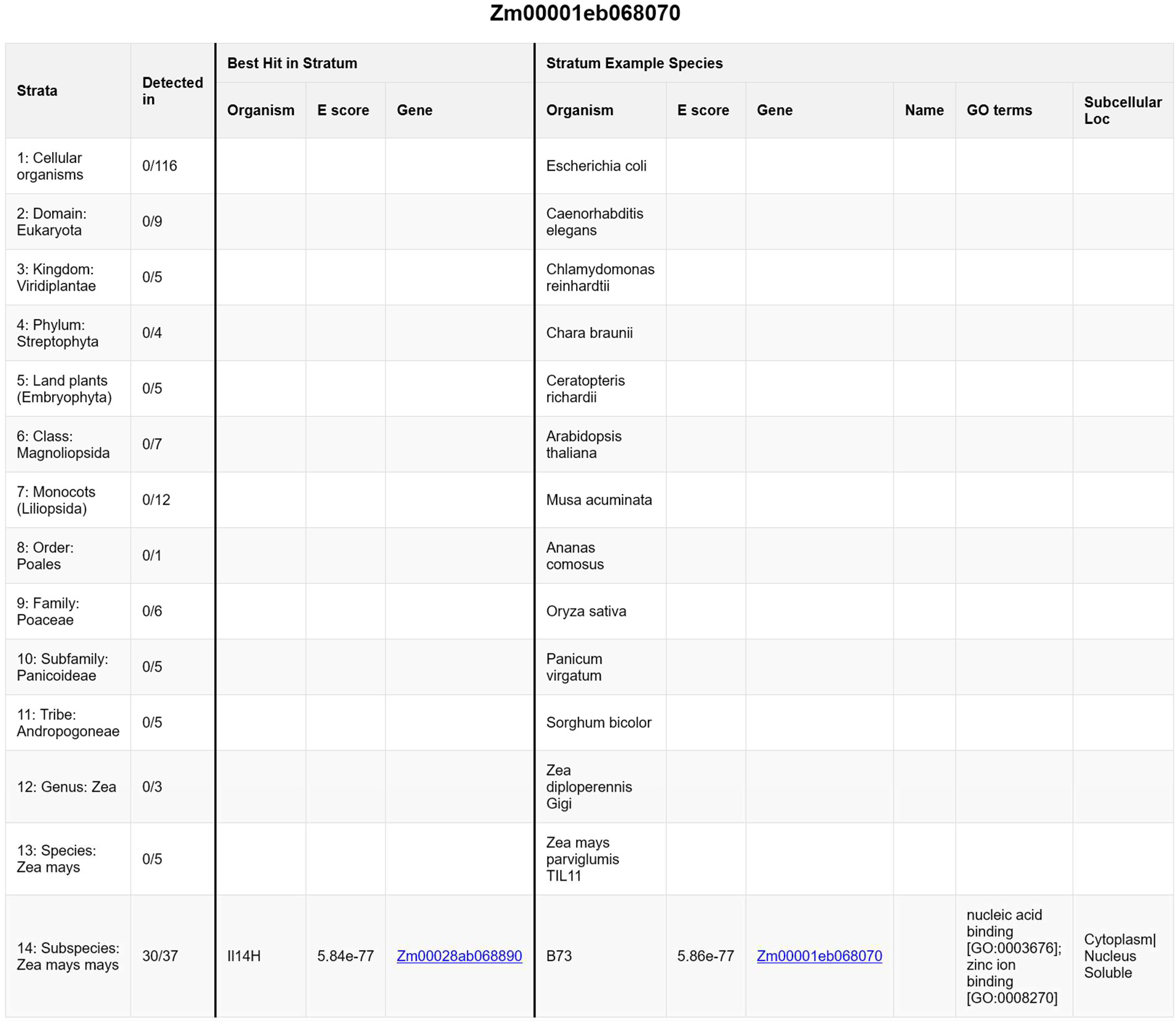
Example Details page, for Zm00001eb068070. This Details page for a maize-specific gene shows GO terms and subcellular localization only for B73, as homologs are not present in other species.

**Fig. S4:**
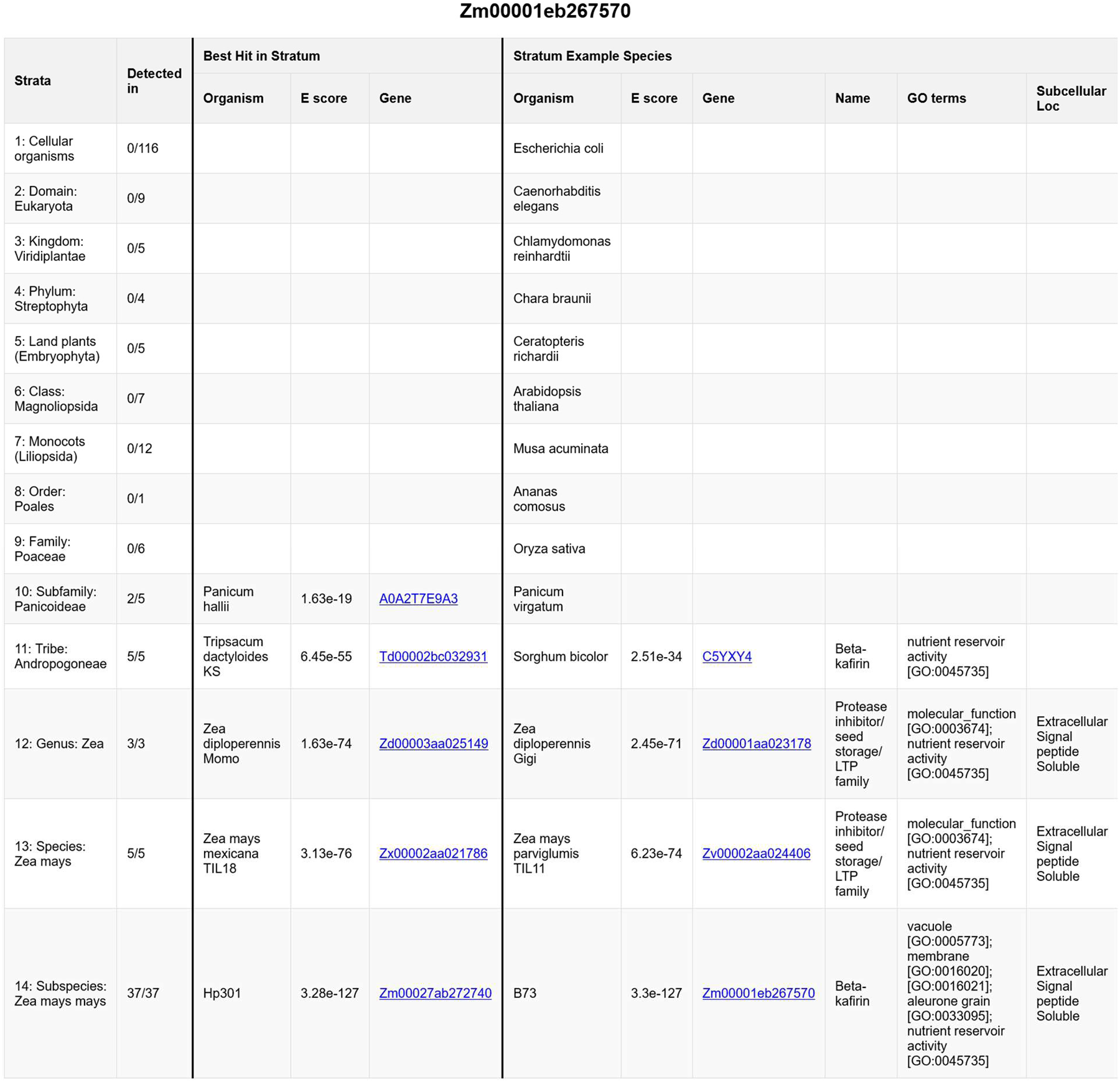
Example Details page, for Zm00001eb267570 (zp15). This Details page shows similar GO terms and subcellular localization across example species, indicating likely conservation of function throughout the subfamily Panicoideae.

**Fig. S5:**
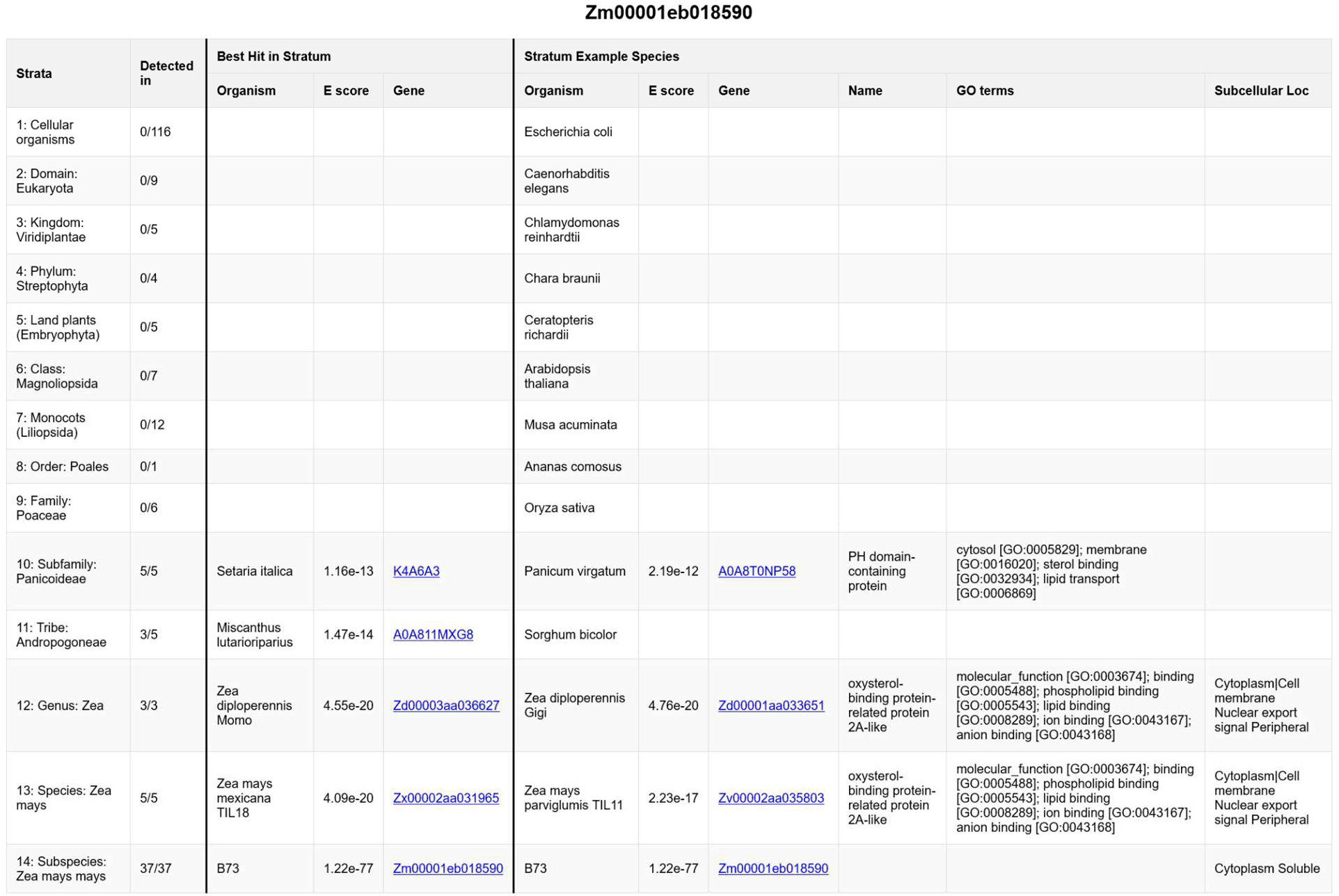
Example Details page, for Zm00001eb018590.

**Fig. S6:**
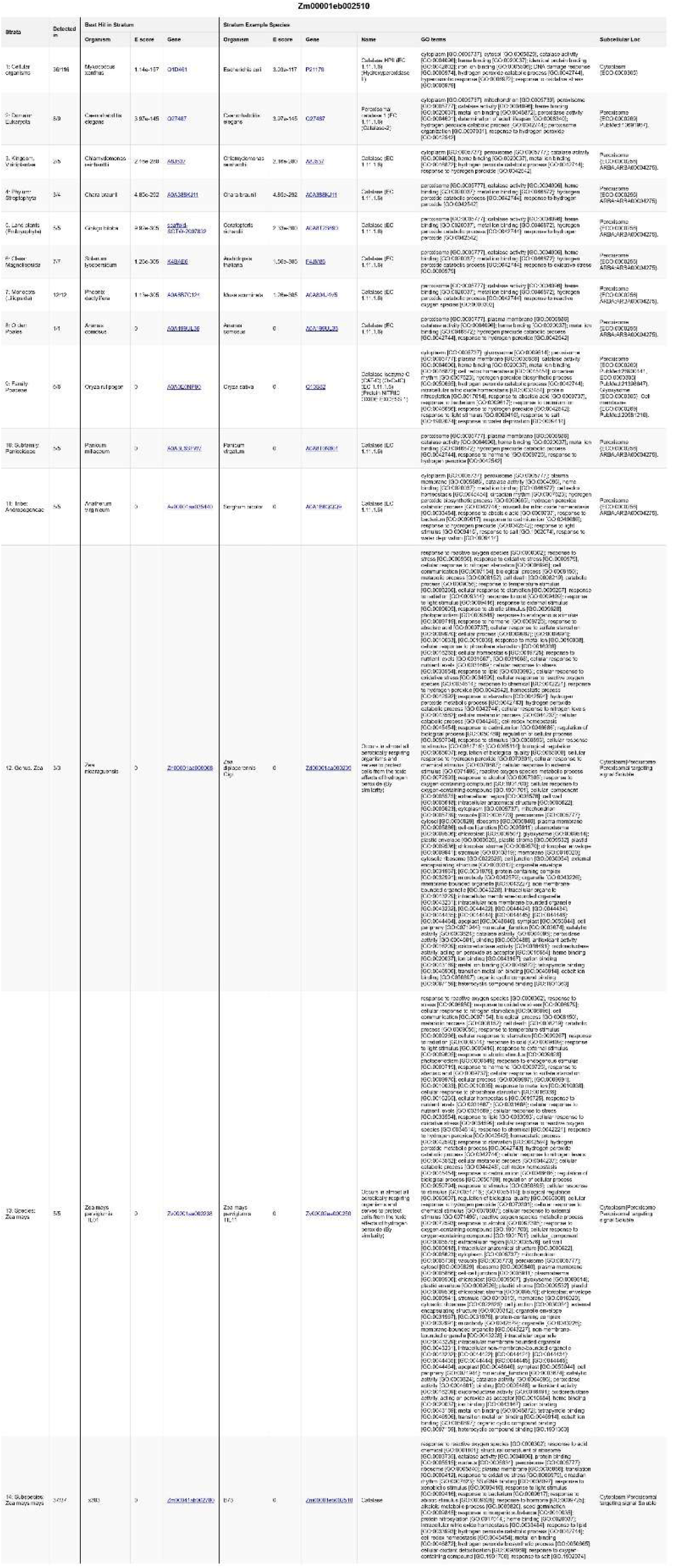
Full Details page for Zm00001eb002510 (cat2) (Fig. 1C).

**Fig. S7:**
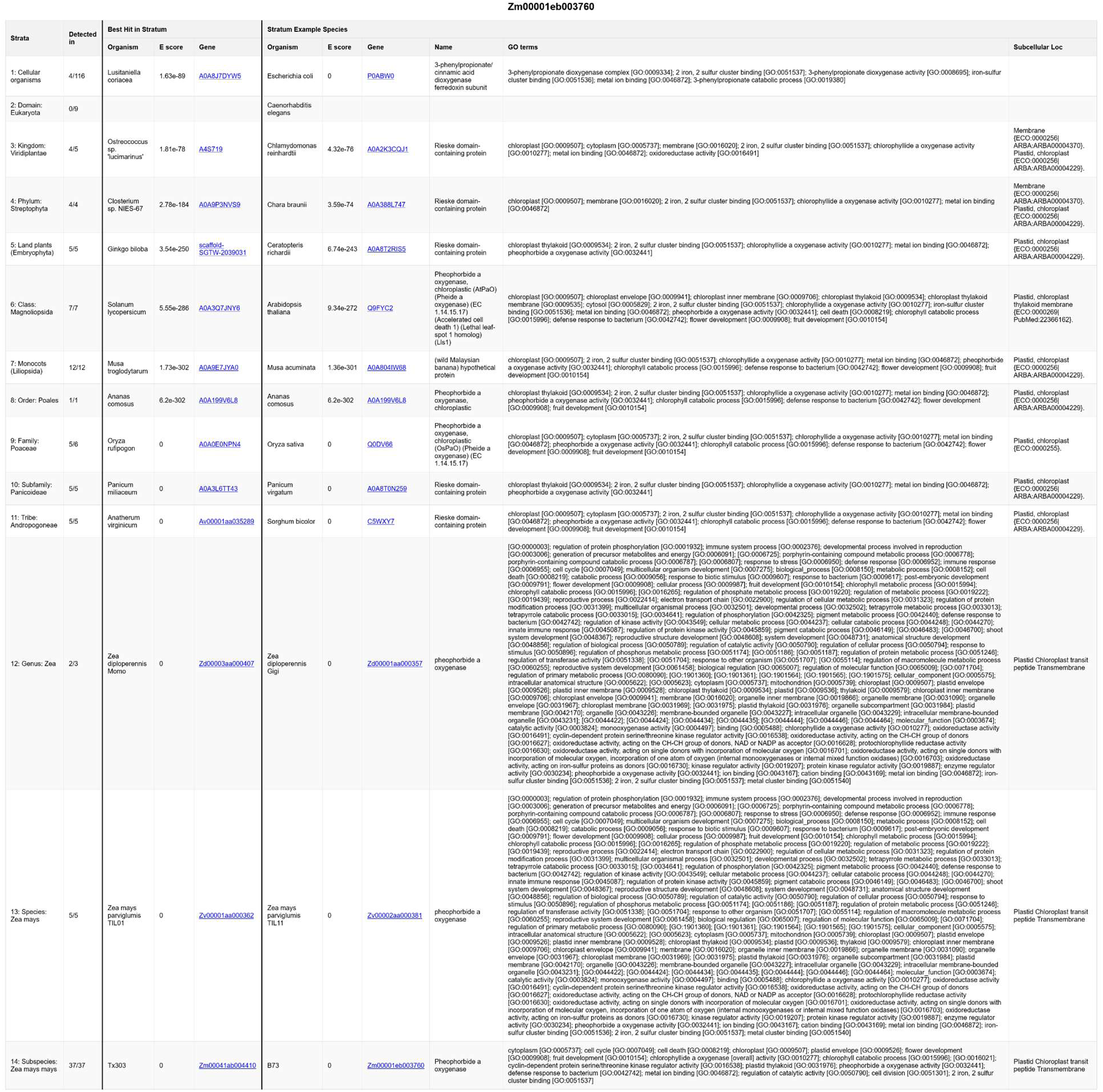
Example Details page, for Zm00001eb003760 (lls1). This Details page shows the conserved function of lls1 homologs across all cellular organisms as an oxygenase, but also the GO term defense response to bacterium [GO:0042742] beginning in Phylostratum 6 (class: Magnoliopsida) indicating an additional, new function for this conserved protein.

**Fig. S8:**
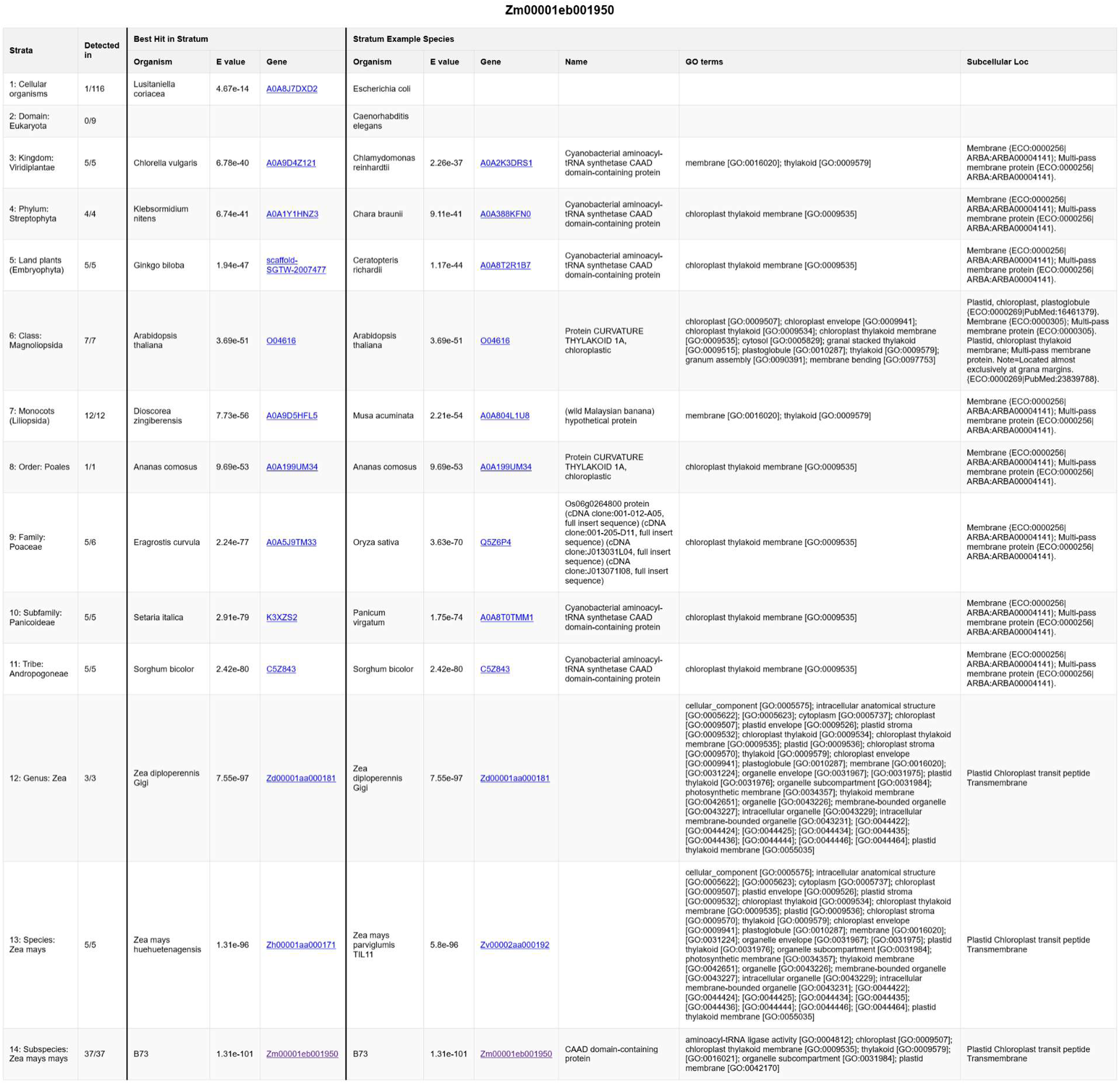
Example Details page, for Zm00001eb001950 (pba1), showing the skipped Phylostratum 2 reflecting horizontal gene transfer.

**Fig. S9:**
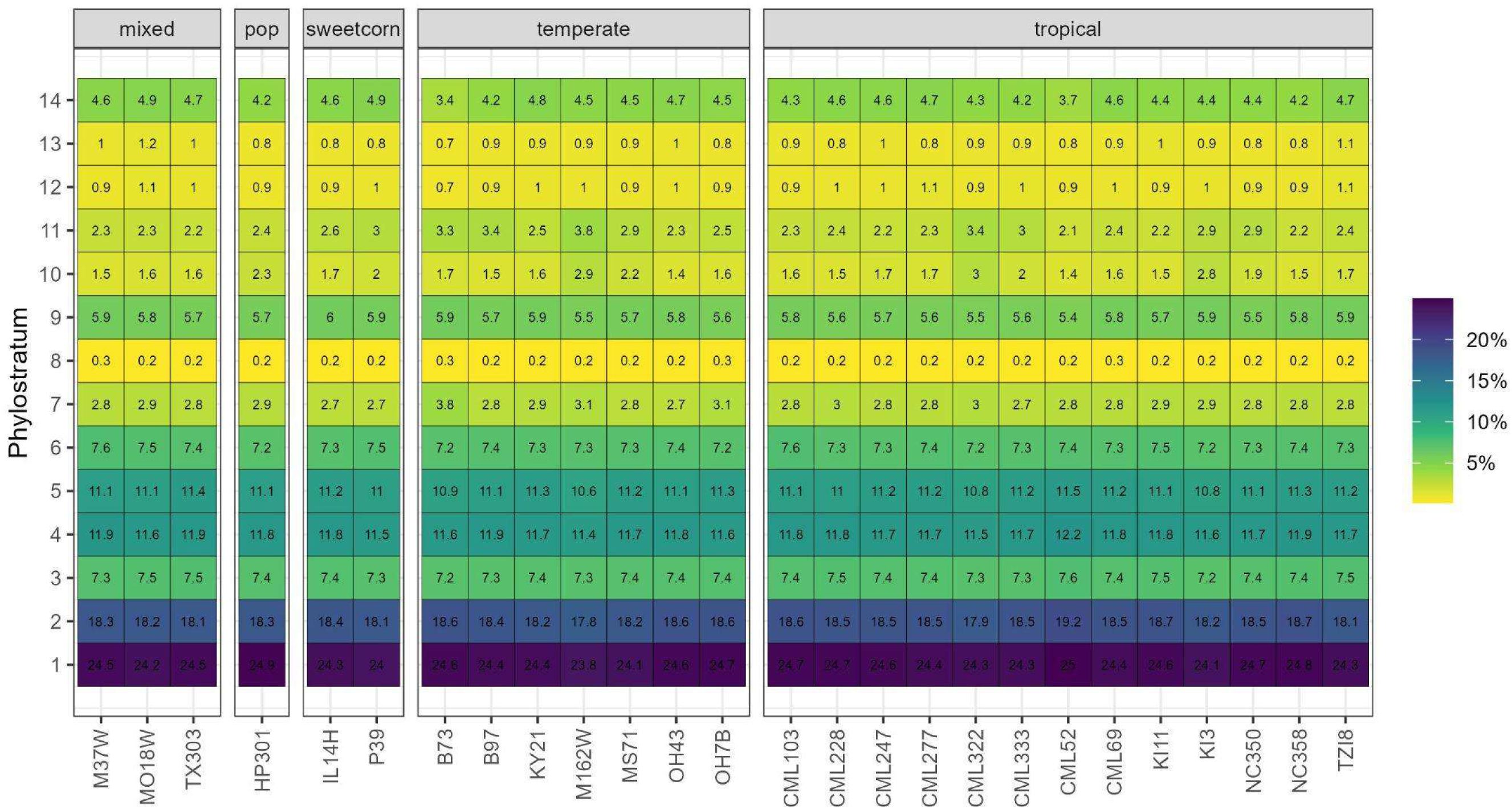
Percentage of proteins in each phylostratum by NAM founder, faceted 528 by germplasm group.

**Fig. S10:**
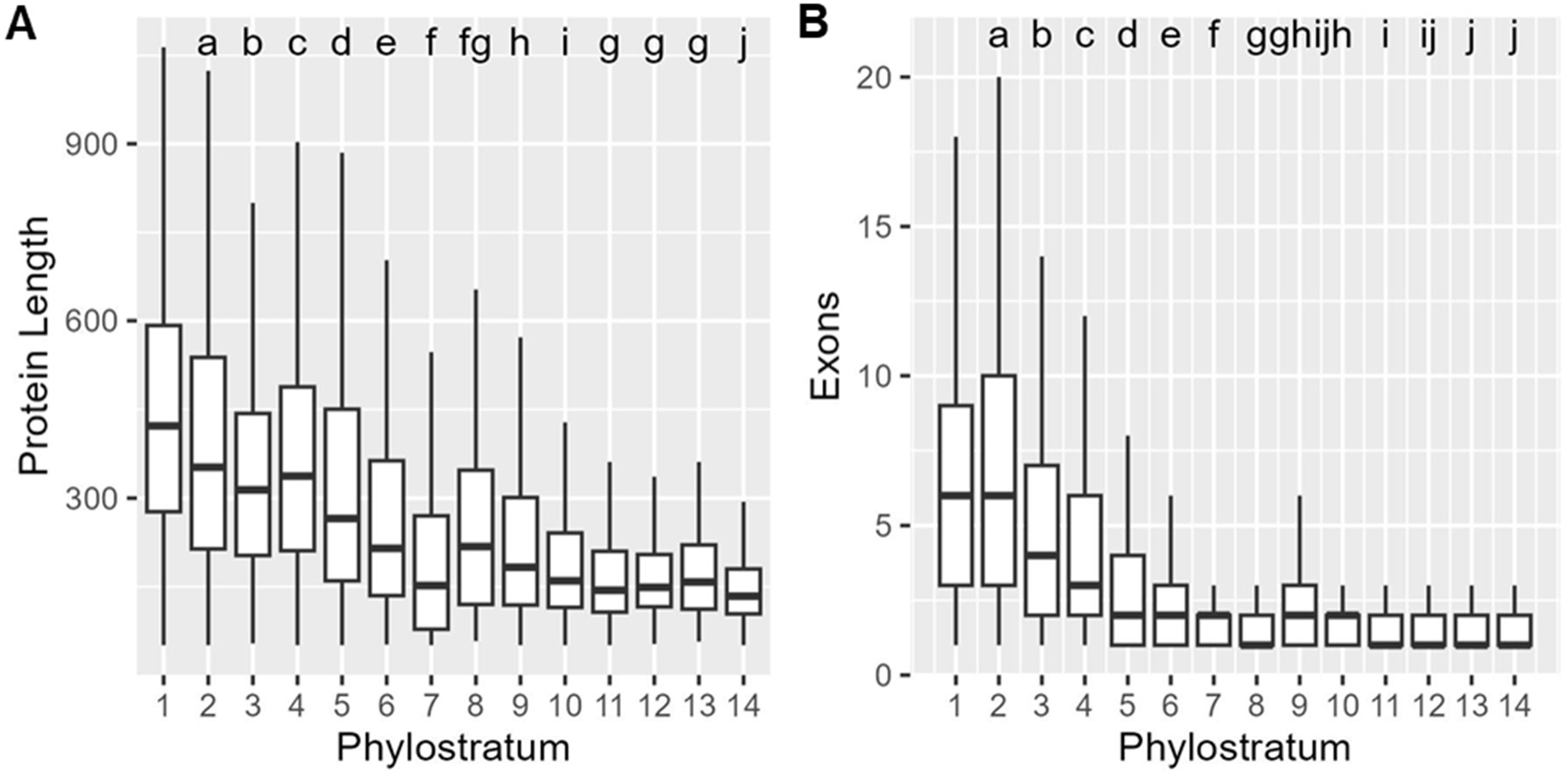
Protein length and exon number by phylostratum in B73. Shared letters indicate phylostrata that are not significantly different using a pairwise Wilcoxon test with FDR correction for multiple testing. Outliers above 1.5 IQR from the median not plotted for clarity. A) Protein length and B) exon number.

**Fig. S11:**
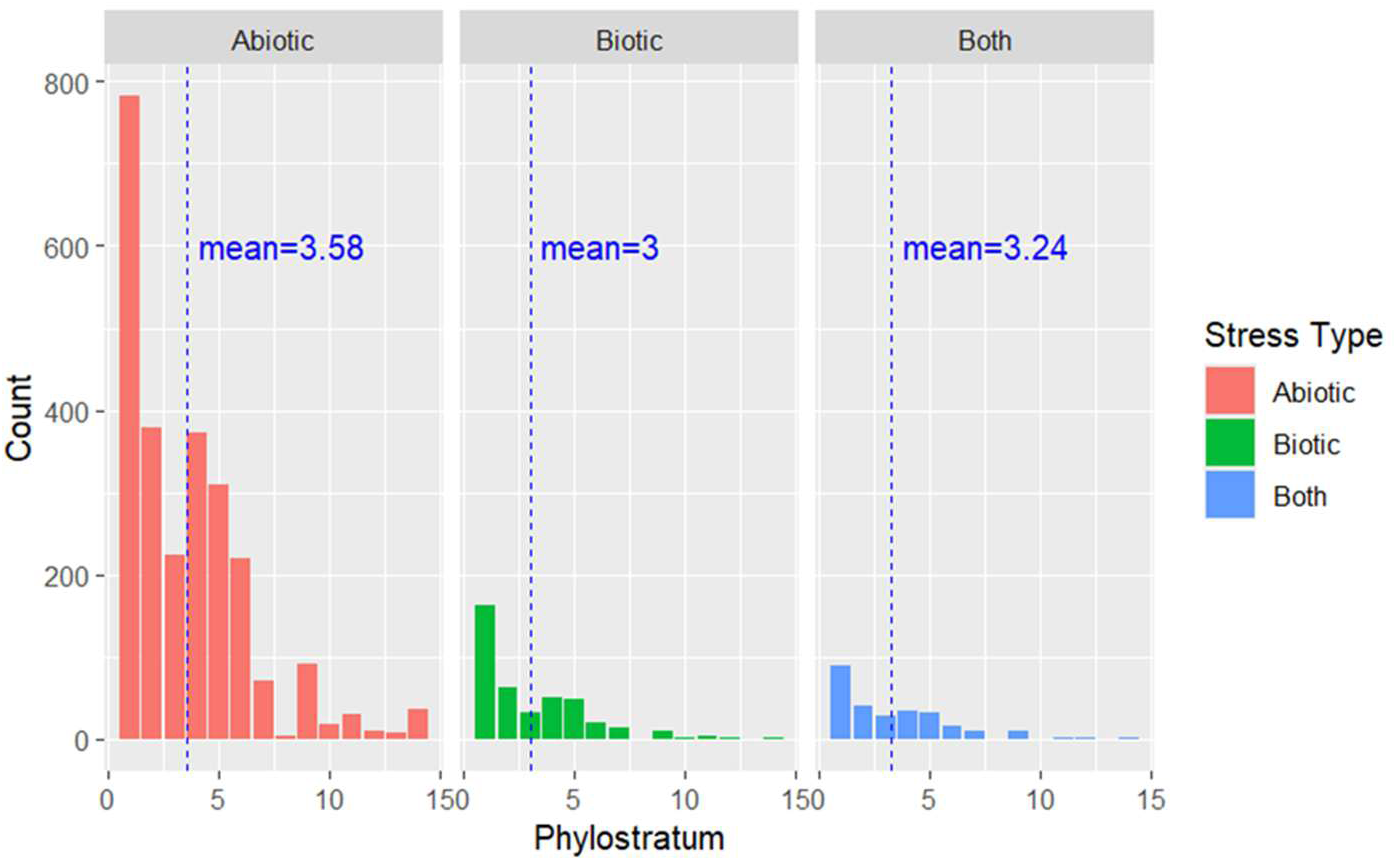
Distribution of phylostrata of B73 DEGs responsive to abiotic, biotic, or both types of stress.

**Fig. S12:**
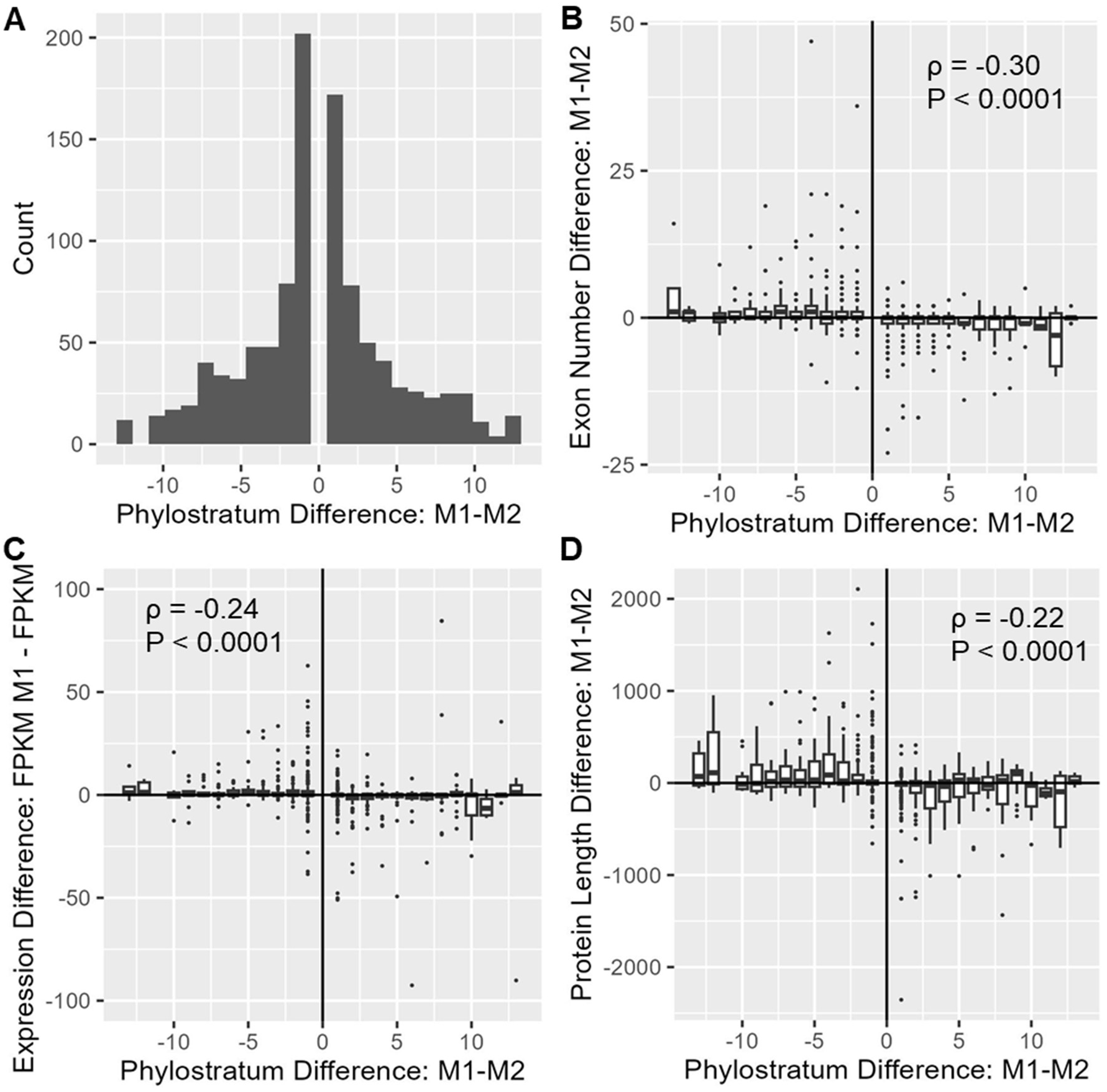
20% of maize homeologs were assigned to different phylostrata in Subgenome 1 (M1) than Subgenome 2 (M2). The left part of each figure (M1-M2<0) shows cases where the M1 homeolog is more conserved and the right (M1-M2>0) shows cases where the M2 homeolog is more conserved. Cases where M1=M2 (M1-M2=0) make up 80% of all homeologous pairs and are not shown. A) Distribution of differences in phylostrata. Differences in B) exon number, C) expression difference, and D) protein length by phylostratum difference.

**Fig. S13:**
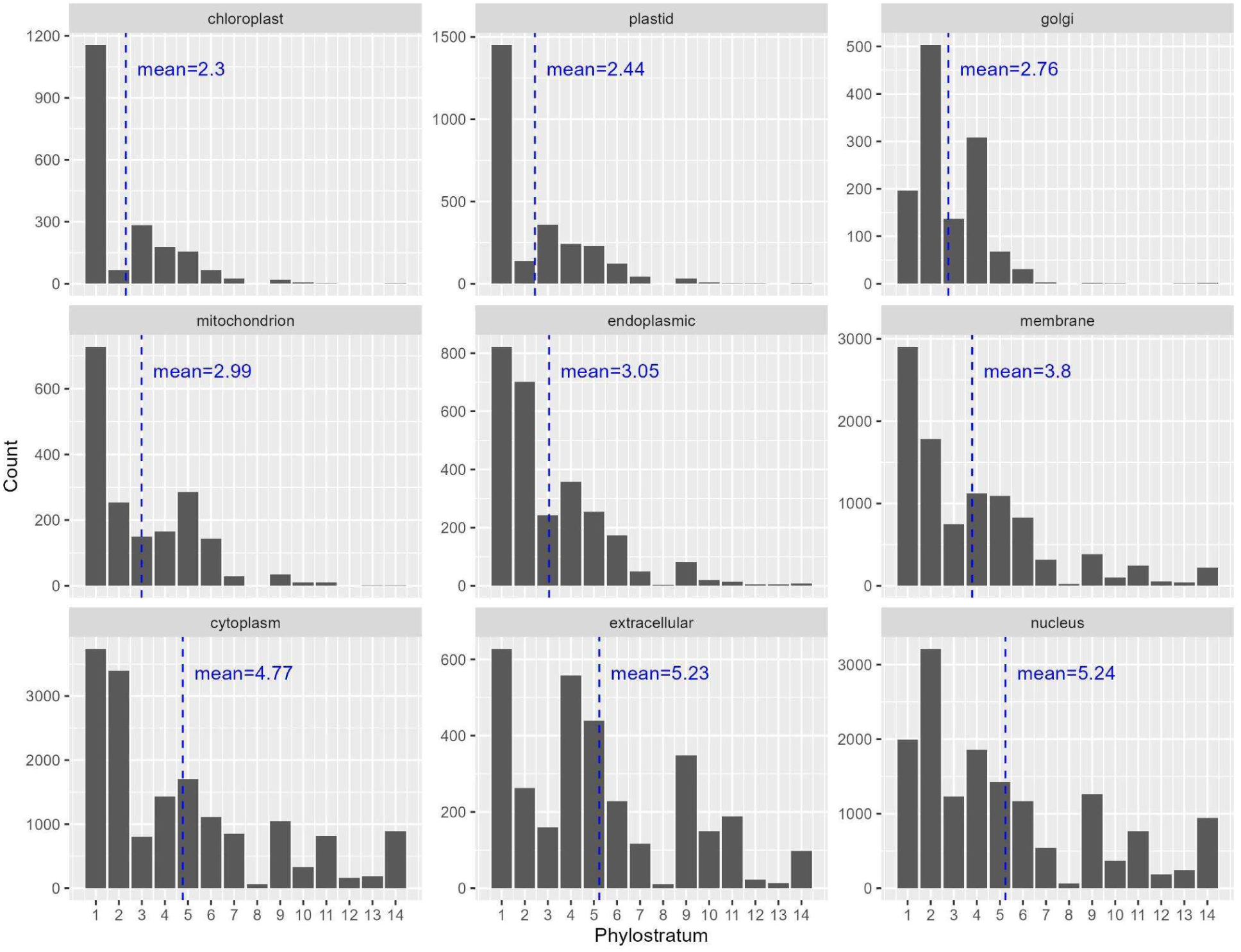
Phylostratum by subcellular localization in B73. Note that some genes were assigned to multiple locations. Subcellular localizations are ordered by mean phylostratum.

**Fig. S14:**
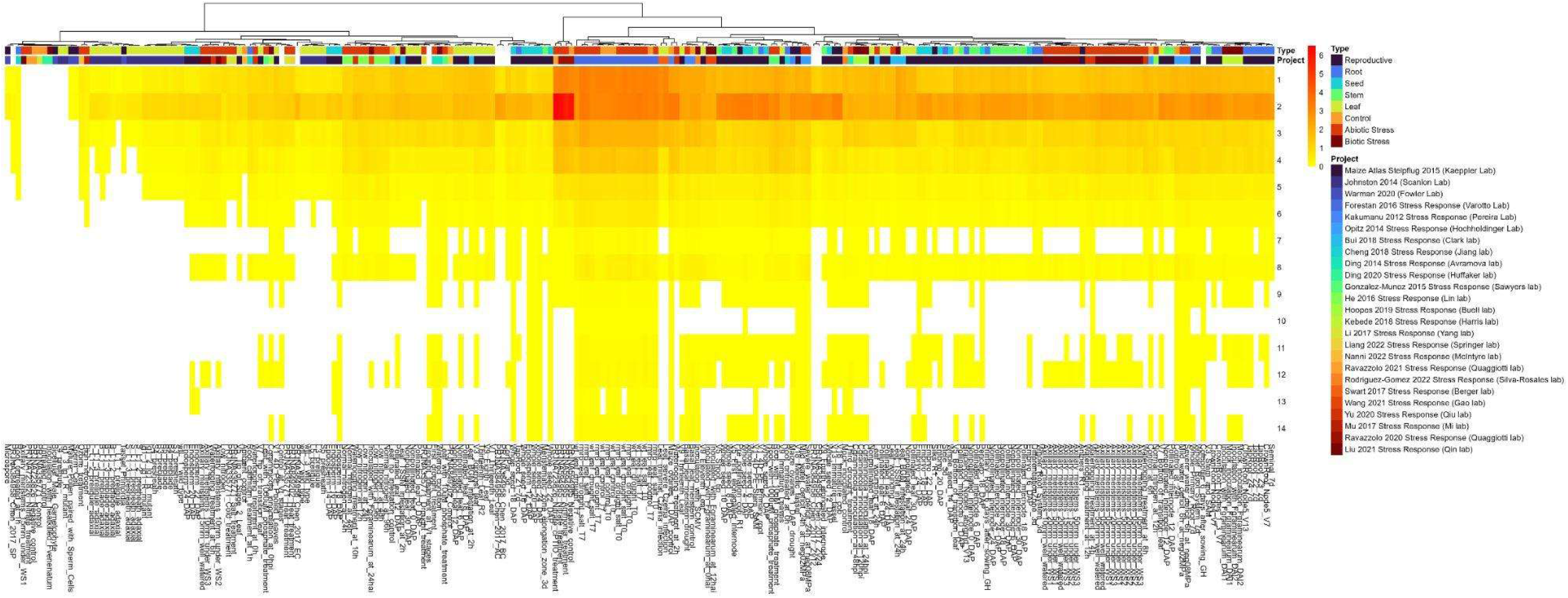
Median expression by phylostratum in 241 B73 tissues. Expression is measured as log_2_ of (FPKM+1), and columns were sorted automatically by ‘pheatmap()’ in R. The tissue type (first row) and project (second row) are shown by color-coded labels at the top of the heatmap.

**Table S1: GO term enrichment by phylostratum in B73.**

